# Comprehensive Transcriptomic Analysis of COVID-19 Blood, Lung, and Airway

**DOI:** 10.1101/2020.05.28.121889

**Authors:** Andrea R. Daamen, Prathyusha Bachali, Katherine A. Owen, Kathryn M. Kingsmore, Erika L. Hubbard, Adam C. Labonte, Robert Robl, Sneha Shrotri, Amrie C. Grammer, Peter E. Lipsky

## Abstract

**Abstract:** SARS-CoV2 is a previously uncharacterized coronavirus and causative agent of the COVID-19 pandemic. The host response to SARS-CoV2 has not yet been fully delineated, hampering a precise approach to therapy. To address this, we carried out a comprehensive analysis of gene expression data from the blood, lung, and airway of COVID-19 patients. Our results indicate that COVID-19 pathogenesis is driven by populations of myeloid-lineage cells with highly inflammatory but distinct transcriptional signatures in each compartment. The relative absence of cytotoxic cells in the lung suggests a model in which delayed clearance of the virus may permit exaggerated myeloid cell activation that contributes to disease pathogenesis by the production of inflammatory mediators. The gene expression profiles also identify potential therapeutic targets that could be modified with available drugs. The data suggest that transcriptomic profiling can provide an understanding of the pathogenesis of COVID-19 in individual patients.

**Graphical Abstract:** 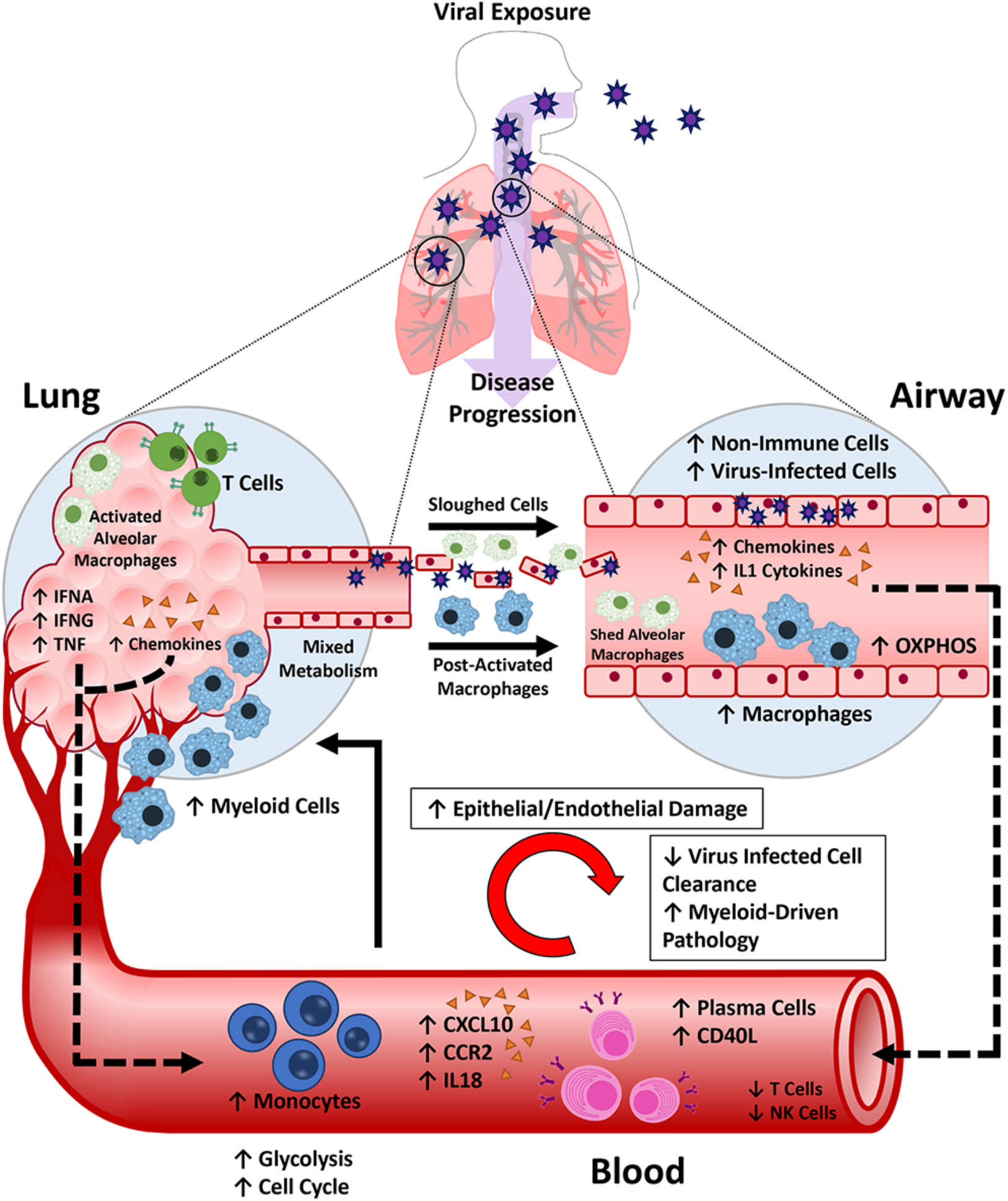

## Introduction

Coronaviruses (CoV) are a group of enveloped, single, positive-stranded RNA viruses causing mild to severe respiratory illnesses in humans (Cui et al., 2019; Fung and Liu, 2019; Greenberg, 2016). In the past two decades, three worldwide outbreaks have originated from CoVs capable of infecting the lower respiratory tract, resulting in heightened pathogenicity and high mortality rates. We are currently in the midst of a pandemic stemming from a third CoV strain, severe acute respiratory syndrome coronavirus 2 (SARS-CoV2), the causative agent of coronavirus disease 2019 (COVID-19). In the majority of cases, patients exhibit mild symptoms, whereas in more severe cases, patients may develop severe lung injury and death from respiratory failure (Chen et al., 2020a; Zhang et al., 2020).

At this time, there is still incomplete information available regarding the host response to SARS-CoV2 infection and the perturbations resulting in a severe outcome. Despite this, clues can be derived from our knowledge of the immune response to infection by other respiratory viruses, including other CoVs. After infection, viruses are typically detected by pattern recognition receptors (PRRs) such as the inflammasome sensor NLRP3, which signal the release of interferons (IFNs) and inflammatory cytokines including the IL-1 family, IL-6, and TNF, that activate a local and systemic response to infection (Kelley et al., 2019; Lazear et al., 2019). This involves the recruitment, activation, and differentiation of innate and adaptive immune cells, including neutrophils, inflammatory myeloid cells, CD8 T cells, and natural killer (NK) cells (Newton et al., 2016). Resolution of infection is largely dependent on the cytotoxic activity of CD8 T cells and NK cells, which enable clearance of virus-infected cells (Newton et al., 2016). Failure to clear virus-infected cells may facilitate a hyper-inflammatory state termed Macrophage (MΦ) activation syndrome (MAS) or “cytokine storm”, and ultimately damage to the infected lung (Crayne et al., 2019; McGonagle et al., 2020).

The recent emergence of SARS-CoV2 and the relative lack of comprehensive knowledge regarding the progression of COVID-19 disease has constrained our ability to develop effective treatments for infected patients. One means to obtain a more complete understanding of the host response to SARS-CoV2 is to examine gene expression in relevant tissues. A limited number of gene expression profiles are available from patients with COVID-19 and have yielded some insights into the pathogenic processes triggered by infection with SARS-CoV2 (Blanco-Melo et al., 2020; Wei et al., 2020; Xiong et al., 2020). However, because of the small number of samples and limited analysis, a full picture of the biological state of SARS-CoV2-affected tissues has not emerged. To address this, we have assessed available SARS-CoV2 gene expression datasets from blood, lung, and airway using a number of orthogonal bioinformatic tools to provide a more complete view of the nature of the COVID-19 inflammatory response and the potential points of therapeutic intervention.

## Results

### Gene expression analysis of blood, lung, and airway of COVID-19 patients

To characterize the pathologic response to SARS-CoV2 infection, we analyzed transcriptomic data from peripheral blood mononuclear cells (PBMCs) and postmortem lung tissue of COVID-19 patients and healthy controls as well as bronchoalveolar lavage (BAL) fluid of COVID-19 patients (CRA002390, GSE147507, **Table S1**) (Blanco-Melo et al., 2020; Xiong et al., 2020). We first determined changes in gene expression in the blood (PBMC-CTL vs PBMC-CoV2) and lung (Lung-CTL vs Lung-CoV2). Because no control BAL fluids were associated with the BAL-CoV2 samples, we compared BAL-CoV2 to PBMC-CoV2 from the same dataset to avoid effects related to batch and methodology. Overall, we found 4,245 differentially expressed genes (DEGs) in the blood (2,166 up and 2,079 down), 2,220 DEGs in the lung (684 up and 1,536 down), and 8,952 DEGs in the airway (BAL) (4,052 up and 4,900 down) (**Table S2**).

### Conserved and differential enrichment of inflammatory cells and pathways in COVID-19 patients

To interrogate pathologic pathways in the 3 compartments, we carried out Gene Set Variation Analysis (GSVA) utilizing a number of informative gene modules (Catalina et al., 2019), (Catalina et al. submitted manuscript) (**Figure 1, Table S3**). Numerous innate immune response pathways were increased in all 3 compartments, whereas adaptive immune signatures tended to be decreased in blood and airway, but not lung, a pattern that is consistent with earlier studies (Blanco-Melo et al., 2020; Wen et al., 2020; Xiong et al., 2020). Although heterogeneity was observed in all compartments, closer examination revealed consistent perturbations. In the blood, the classical and lectin-induced complement pathways, as well as the NLRP3 inflammasome, plasma cells (PCs), and monocytes (Mo) were significantly enriched in COVID-19 patients, whereas cytotoxic cells and neutrophils were decreased. In the lung, the NLRP3 inflammasome, Mo, myeloid cells and low-density granulocytes (LDGs) were enriched in COVID-19 patients, whereas in the airway, the classical and alternative complement pathways were enriched and T cells and cytotoxic cells were decreased.

**Figure 1.**
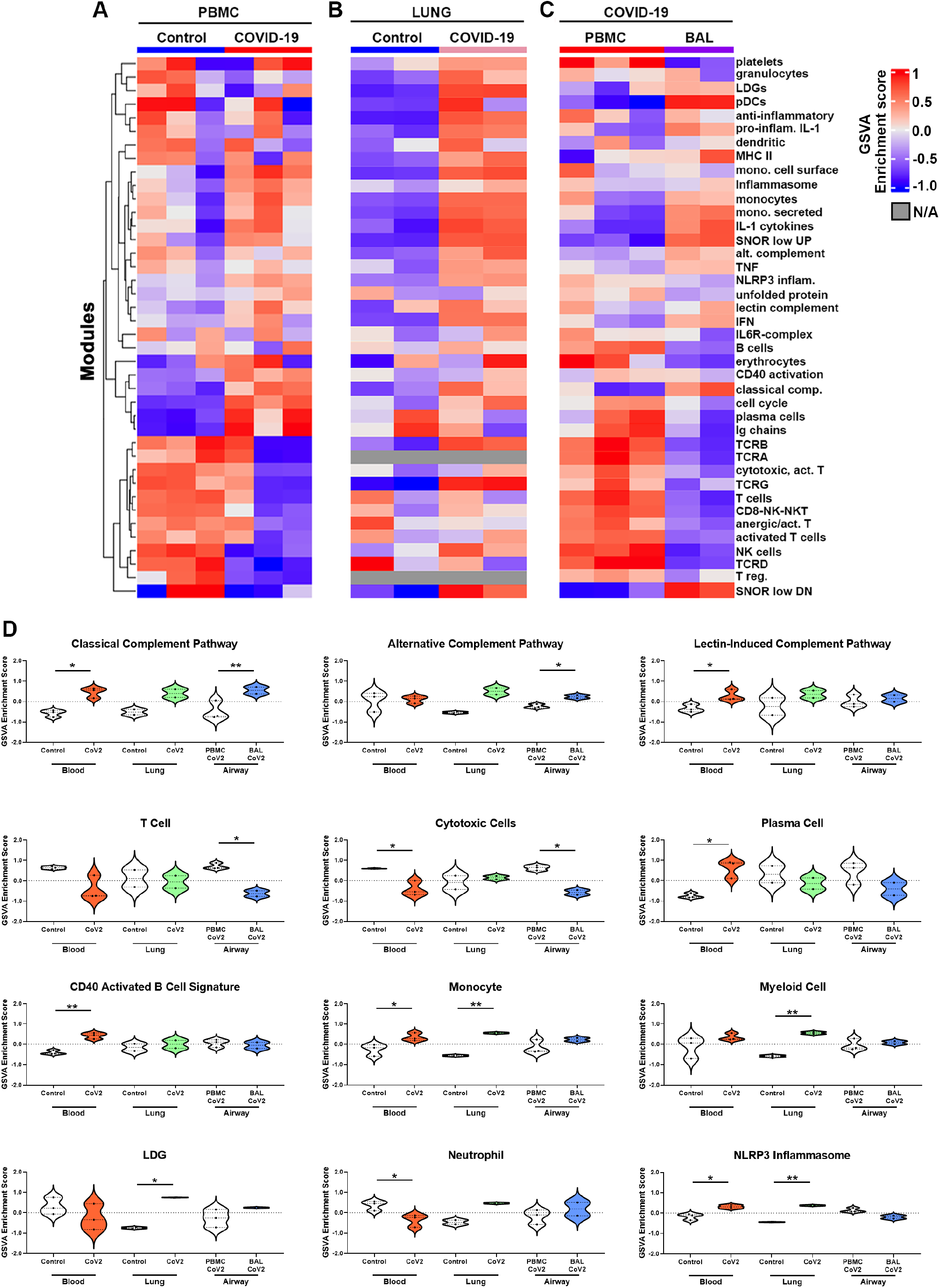
Conserved and differential enrichment of immune cells and pathways in blood, lung, and airway of SARS-CoV2-infected patients. (**A-D**) Individual sample gene expression from the blood (**A**), lung (**B**), and airway (**C**) was analyzed by GSVA for enrichment of immune cell and inflammatory pathways. Select enrichment scores are shown as violin plots in (**D**). *p<0.05, **p<0.01

Although there is conflicting data on the presence of an IFN gene signature (IGS) and whether SARS-CoV2 infection induces a robust IFN response (Blanco-Melo et al., 2020; Trouillet-Assant et al., 2020; Wei et al., 2020), we observed increased expression of Type I IFN genes (*IFNA4, IFNA6, IFNA10*) and significant enrichment of the common Type I and Type II IGS, including enrichment of *IFNA2*, *IFNB1 and IFNG* gene signatures specifically in the lung tissue (**Figure 2A&B**). Furthermore, we detected increased expression of genes found to be important for the anti-viral innate immune response (*IFIH1, DDX58, EIF2AK2, OAS2*) and decreased expression of negative regulators of this response (*IRF2BP1, SKIV2L*) in both the lung and airway compartments (**Figure 2C**) (Fischer et al., 2020). Interestingly, we found expression of *MAVS*, a signaling adaptor for RNA virus sensors, was decreased in the airway, which is consistent with the reported effect of SARS-CoV2 and may reflect a mechanism of viral immune evasion (Bojkova et al., 2020; Vazquez and Horner, 2015).

**Figure 2.**
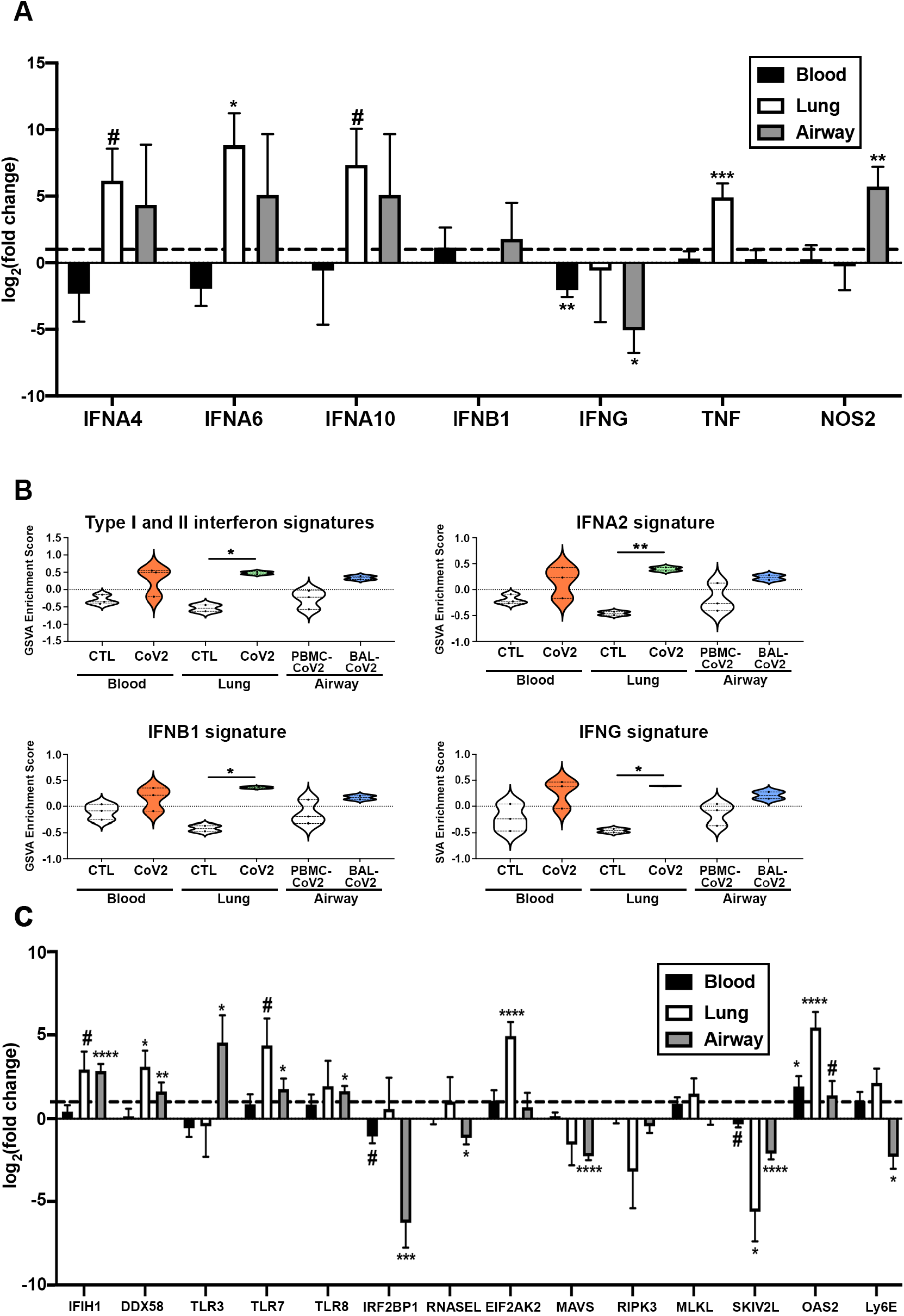
Elevated IFN expression in the lung tissue of COVID-19 patients. **(A)** Normalized log2 fold change RNA-seq expression values for IFN-associated genes from blood, lung, and airway of individual COVID-19 patients. The dotted line represents the expression of each gene in healthy individuals (for blood and lung) or PBMCs from COVID-19 patients (airway). **(B)** Individual sample gene expression from the blood, lung, and airway was analyzed by GSVA for enrichment of IFN-related gene signatures. (**C**) Normalized log2 fold change RNA-seq expression values for anti-viral genes as in (**A**). #p<0.2, ##p<0.1, *p<0.05, **p<0.01, ***p<0.001, ****p<0.0001

### Increased expression of inflammatory mediators in the lungs of COVID-19 patients

To examine the nature of the inflammatory response in the tissue compartments in greater detail, we examined specific DEGs of interest (**Figure 3A&B**). In the blood, we noted increased expression of the inflammatory chemokine *CXCL10*, which is an IFNG response gene and involved in the activation and chemotaxis of peripheral immune cells, (Lindell et al., 2008), the chemokine receptor *CCR2*, which has been shown to be critical for immune cell recruitment in response to respiratory viral infection (Teijaro, 2015), as well as the inflammatory IL-1 family member, *IL18*. Expression of a number of chemokines, including ligands for CCR2, were significantly increased in both the lung tissue and airway of COVID-19 patients, including *CCL2, CCL3L1, CCL7, CCL8,* and *CXCL10*. We also observed elevated pro-inflammatory IL-1 family members, *IL1A* and *IL1B,* in these 2 compartments. Furthermore, lung tissue exhibited enrichment of the IL-1 cytokine gene signature, whereas the airway exhibited additional expression of *IL18, IL33, IL36B,* and *IL36G*.

**Figure 3.**
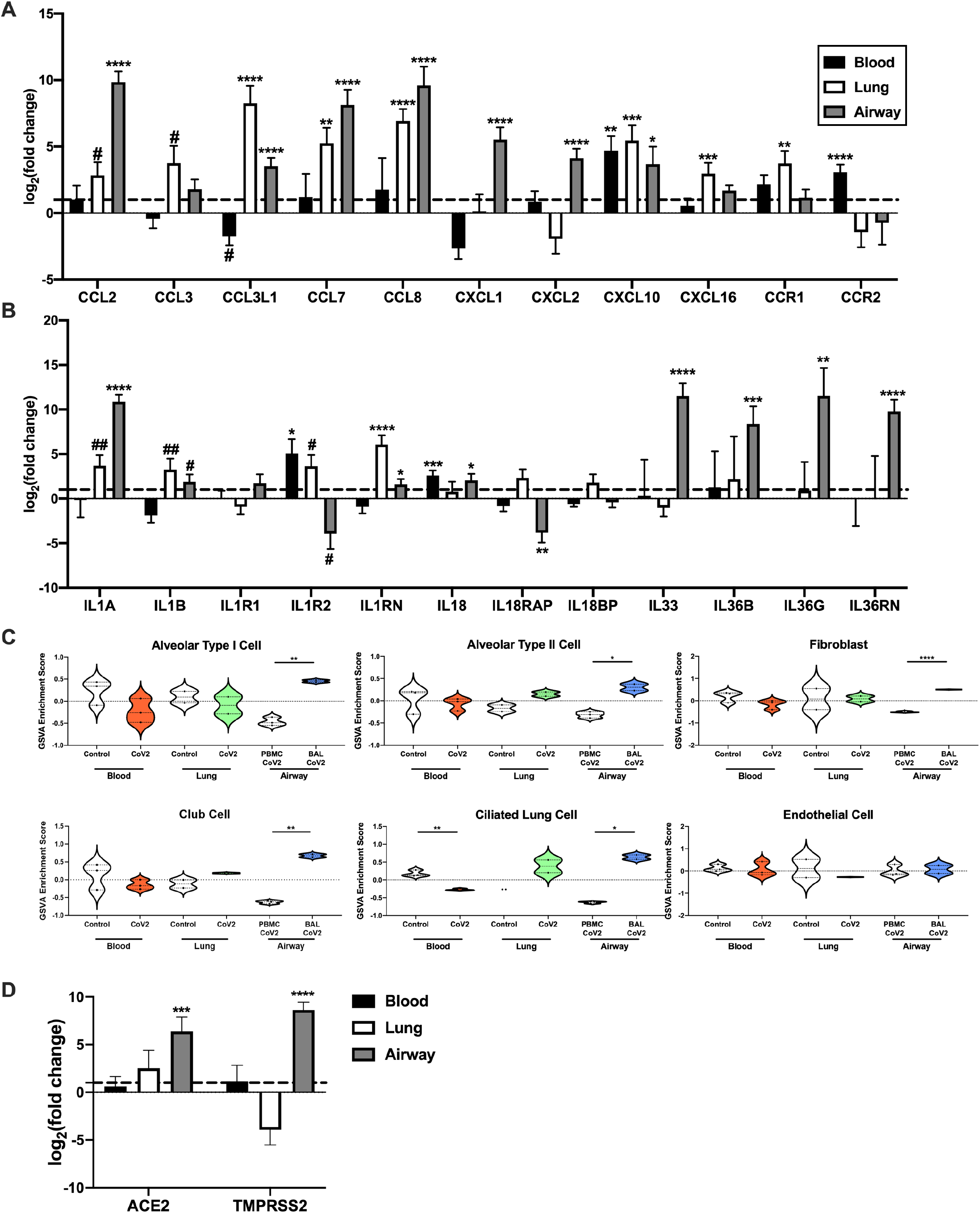
Viral entry gene expression correlates with enhanced expression of inflammatory mediators in SARS-CoV2-infected lungs. (**A-B**) Normalized log_2_ fold change RNA-seq expression values for chemokines and chemokine receptors (**A**) and IL-1 family members (**B**) from blood, lung, and airway of COVID-19 patients as in **Figure 3A**. (**C**) Individual sample gene expression from the blood, lung, and airway was analyzed by GSVA for enrichment of various lung tissue cell categories. (**D**) Normalized log_2_ fold change RNA-seq expression values for viral entry genes as in (**A&B**). #p<0.2, ##p<0.1, *p<0.05, **p<0.01, ***p<0.001, ****p<0.0001

### Non-hematopoietic cells in the BAL fluid may be indicative of viral-induced damage

To determine whether viral infection resulted in modification of resident tissue populations, we employed GSVA with various non-hematopoietic cell gene signatures (**Figure 3C**). We found that signatures of various lung tissue cells but not endothelial cells were enriched in the airway, but not the lung of COVID-19 subjects. Additionally, we also detected increased expression of the viral entry genes *ACE2* and *TMPRSS2*, which are typically expressed on lung epithelium (Sungnak et al., 2020) (**Figure 3D**).

### Protein-protein interactions identify myeloid subsets in COVID-19 patients

We next sought to utilize an unbiased, protein-protein interaction (PPI)-based clustering approach to assess the inflammatory cell types within each tissue compartment. PPI networks predicted from DEGs were simplified into metastructures defined by the number of genes in each cluster, the number of significant intra-cluster connections, and the number of associations connecting members of different clusters to each other (**Figure 4A-C, Table S4**). Overall, upregulated PPI networks identified numerous specific cell types and functions. In the blood, cluster 8 was dominated by a Mo population expressing *C2*, *C5*, *CXCL10*, *CCR2,* and multiple IFN-stimulated genes, whereas cluster 3 contained hallmarks of alternatively activated (M2) macrophages (MΦs) and/or myeloid-derived suppressor cells (MDSCs), including *CD33*, *CD36*, *CD93,* and *ITGAM* (**Figure 4A).** Smaller immune clusters were indicative of functions, including inflammasome activation, damage-associated molecular patterns (DAMP) activity, the classical complement cascade and the response to Type II IFNs. Myeloid heterogeneity in the blood was also reflected by the presence of multiple metabolic pathways, such as enhanced oxidative phosphorylation (OXPHOS) in cluster 1 linked to M2-like MΦs in cluster 3 (mean interaction score of 0.875), and glycolysis in clusters 7 and 13 connected to activated Mo in cluster 8 (interaction scores of 0.86 and 0.82, respectively). Consistent with our GSVA results, blood exhibited profoundly decreased T cells determined by the downregulation of T cell activation markers *CD28*, *LCK* and *ITK* (**Figure S1A**).

**Figure 4.**
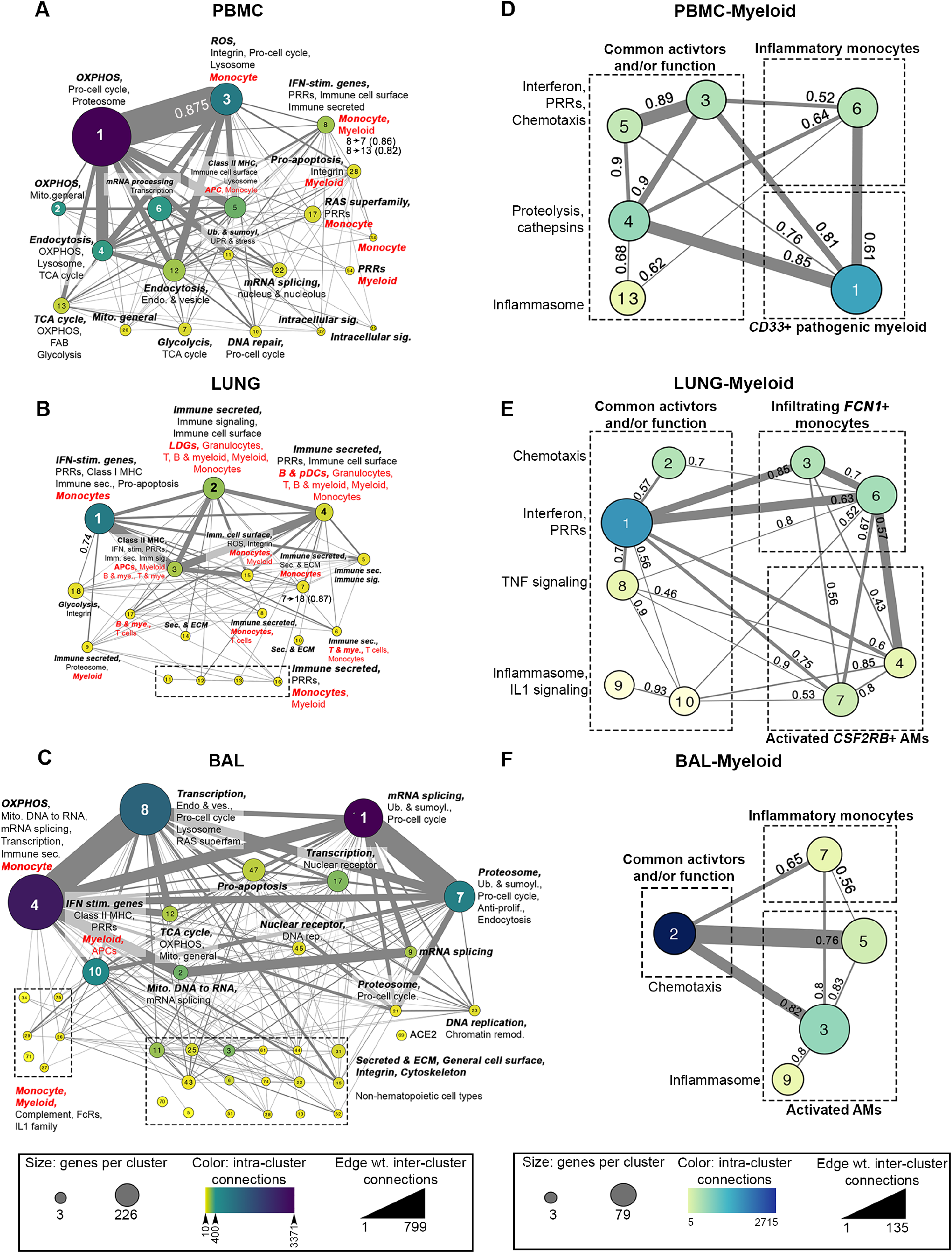
PPI analysis identifies different myeloid cell subsets and metabolic pathways in blood, lung, and airway of COVID-19 patients. DE upregulated genes from blood (**A**), lung (**B**) and airway (**C**) were used to create PPI metaclusters. Size indicates the number of genes per cluster, color indicates the number of intra-cluster connections and edge weight indicates the number of inter-cluster connections. Enrichment for biological function and immune cell type was determined by BIG-C and I-Scope, respectively. Small clusters (~14 genes) with similar function are grouped in dotted-line boxes. Clusters enriched in Mo/myeloid genes were combined by decreasing cluster stringency to create a new set of myeloid-derived metastructures from the blood (**D**), lung (**E**) and airway (**F**). Interaction scores showing the strength of interaction between clusters are indicated (0.4-0.6, medium interaction; 0.61-0.8, strong interaction; 0.81-0.99, very strong interaction).

In addition to the various Mo/myeloid populations, lung tissue was infiltrated by LDGs, granulocytes, T cells, and B cells. Metabolic function in the lung was varied, with Mo-enriched clusters (1 and 7) linked to glycolysis in cluster 18 (interaction scores of 0.74 and 0.87, respectively) potentially reflecting cellular activation, whereas OXPHOS was predominantly downregulated along with other nuclear processes (transcription and mRNA processing) (**Figure 4B, Figure S1B**). The airway was enriched in inflammatory Mo, mitochondrial function and transcription. Multifunctional cluster 4 was dominated by numerous chemokine and cytokine receptor-ligand pairs, whereas smaller immune clusters were enriched in classical complement activation, IFNG and IL-1 responses. (**Figure 4C**). Consistent with tissue damage, we found numerous small clusters in the airway, reflecting the presence of non-hematopoietic cells, including those containing multiple intermediate filament keratin genes, cell-cell adhesion claudin genes and surfactant genes. Notably, non-hematopoietic cell signatures in the airway were similar in content to those derived from *in vitro* SARS-CoV-2-infected primary lung epithelial cell lines (NHBE) (Blanco-Melo et al., 2020) (**Figure S1D**).

Given the large number of clusters including Mo/myeloid/MΦ, we next examined these clusters in greater detail by altering the stringency of PPI clustering to further characterize unique myeloid lineage cells within each tissue compartment (**Figure 4D-F, Table S4**). Myeloid lineage-specific clusters were then compared to previously published gene signatures, including populations G1-G4 reported in BAL of COVID-19 patients (Liao et al., 2020) (**Figure S2A**). In the blood, we found gene modules representative of common myeloid function (chemotaxis, proteolysis, etc.), as well as two independent Mo/myeloid subpopulations (**Figure 4D**). Cluster 6 contained numerous markers highly reminiscent of classically activated blood Mo and exhibited significant overlap with the inflammatory G1 population (Liao et al., 2020), whereas cluster 1 was similar to IFN-activated MΦs, CX3CR1+ synovial lining MΦs (from arthritic mice) and alveolar MΦs (AM) (**Figure S2A**).

In the lung (**Figure 4E**), clusters 2, 3 and 6 overlapped with the G1 inflammatory Mo population and expressed a number of chemotaxis genes. A second population characteristic of AMs was also evident in the lung, defined by *CSF2RB*, the receptor for GM-CSF, a cytokine that regulates AM differentiation (Joshi et al., 2006; Newton et al., 2016; Suzuki et al., 2014). Further characterization of this population indicated significant expression of the coagulation system genes *F5*, *FGG, FGL1*, *SERPINA* and *SERPINE2*. Similarly, re-clustering of Mo/MΦs/myeloid clusters from the airway revealed a population with hallmarks of inflammatory/M1 MΦ (*MARCO* and multiple members of the complement cascade; cluster 7), and a second population of AMs (**Figure 4F**) demonstrating significant overlap with the G3 and G4 populations (**Figure S2A**) (Liao et al., 2020).

### Characterization of myeloid populations in COVID-19 patients

The overlap between previously characterized BAL-defined gene signatures from COVID-19 patients (Liao et al., 2020) and tissue-defined PPI clusters motivated us to evaluate these populations in greater detail by GSVA. Consistent with PPI clusters, the inflammatory-MΦ G1 population was increased in the blood (**Figure S2B**). The G1 and G1 & G2 populations were increased in the lung, consistent with the expression of IFN and pro-inflammatory cytokines (**Figure S2C**). In the airway, the G2, G3, and G4 populations were significantly enriched indicating the presence of both pro-inflammatory MΦs and AMs (**Figure S2D-F**). As a whole, we found that gene signatures of previously defined Mo/MΦ populations in COVID-19 BAL were dispersed among the blood, lung, and airway compartments.

### Co-expression further delineates Mo/MΦ gene expression profiles of COVID-19 patients

We next sought to identify the biology of the populations of Mo/MΦ in the tissue compartments in greater detail. We derived a set of 196 co-expressed Mo/myeloid genes and used them (**Figure S3, Table S5, see Methods**) to probe heterogeneity in each tissue compartment (**Figure 5A**). Notably, we found co-expression of 40 core genes between all compartments, which included complement, chemokine, and cytokine genes (**Figure 5B&C**). In addition, there were 86 shared co-expressed genes in lung and blood, 57 in the lung and airway, and 61 in the airway and blood (**Figure 5B**). To directly compare levels of these 196 co-expressed myeloid genes in each compartment, we normalized gene expression in each sample using 3 genes included in the core 40 genes, (*FCGR1A, FCGR2A, FCGR2C*) (**Figure 5D**). Although many genes were not significantly different between compartments, numerous chemokines and cell surface markers (*CCL2*, *CCL7*, *CCL8*, *CXCL10*, *CLEC4E, FCER1G)* and inflammatory cytokines (*IL1A* and *TNF)* were enriched in the lung compared to the blood and airway. Furthermore, the complement genes *C1QB*, *C1QC*, and *C2* were increased in the lung compared to the blood, but not changed between the lung and the airway. Altogether, these normalized gene expression results suggested that expression of inflammatory mediators was increased in SARS-CoV2-infected lung over the other compartments and in the airway compared to the blood.

**Figure 5.**
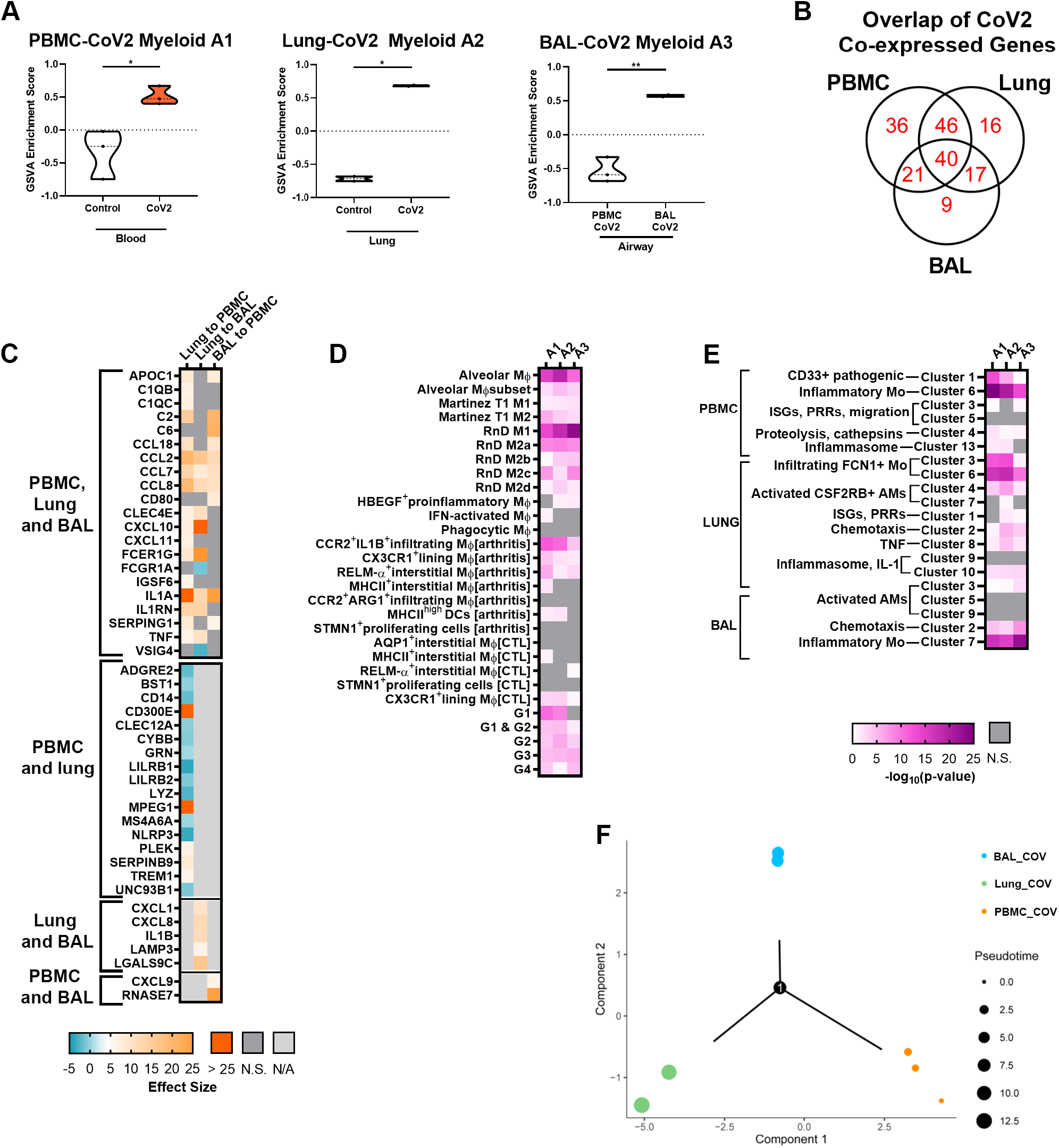
Different co-expression-derived myeloid populations are found in blood, lung, and airway of COVID-19 patients. (**A**) Individual sample gene expression from the blood, lung, and airway was analyzed by GSVA for enrichment of Mo cell surface and secreted categories. (**B**) GSVA enrichment of myeloid subpopulations increased in COVID-19 blood (A1), lung (A2), and airway (A3). (**C**) Venn Diagram of the gene overlap between myeloid subpopulations A1-A3. (**D**) Comparison of normalized log_2_ fold change expression values of genes defining A1-A3. Expression values for each sample in each comparison were normalized by the mean of the log_2_ fold change expression of *FCGR1A*, *FCGR2A*, and *FCGR2C*. Significant comparisons are displayed by Hedge’s G effect size. (**E-F**) Characterization of A1-A3 by enrichment of previously described myeloid populations (**E**) (**Table S3 & S6**) and PBMC, lung, and BAL myeloid metaclusters from **Figure 4D-F** (**F**). Fisher’s Exact Test was used to calculate overlap between transcriptomic signatures and significant overlaps (p<0.05) are shown as the negative logarithm of the p-value. (**G**) Trajectory analysis using expression of 621 genes (196 myeloid-specific genes used in **B&C**+ 425 additional myeloid genes shown in **Table S5**) in the blood, lung, and airway compartments. Colors represent sample identity and size represents pseudotime distance along the trajectory.

To determine the function and nature of these myeloid populations, we compared them to previously published myeloid signatures (**Figure 5E, Table S6**) (Culemann et al., 2019; Kuo et al., 2019; Liao et al., 2020; Martinez et al., 2006; Reyfman et al., 2019). The population increased in the blood (A1) was predominantly characterized by features of AMs, M1 and M2 MΦs, pro-inflammatory MΦs with potential to infiltrate tissue, and the inflammatory MΦ G1 population. The A1 population also exhibited features of inflamed murine residential, interstitial MΦs. The myeloid cell population increased in COVID lung (A2) was most similar to pro-fibrotic AMs, M1 MΦs, M2 MΦs, blood-derived infiltrating MΦs, and the inflammatory Mo G1 population. A2 was also marked by additional AM-specific genes, contributing to the observed overlap with the other two compartments. However, overlap between A2 and the G4 AM signature was relatively decreased, suggesting that the lung AMs are more similar to those found in pulmonary fibrosis (Reyfman et al., 2019). Finally, the population increased in the airway (A3) similarly exhibited characteristics of AMs, M1 and M2 MΦs, and pro-inflammatory MΦs that have infiltrated into the tissue compartment (**Figure 5E**). Of note, the airway A3 population was not similar to the previously described BAL-derived inflammatory MΦ G1 population (Liao et al., 2020).

We also evaluated the overlap between the Mo/MΦ A1-A3 gene clusters and those identified using PPI clustering (Figure 4) (**Figure 5F, Table S6**). Interestingly, the CD33^+^ pathogenic population (PPI-derived PBMC Myeloid Cluster 1) was most strongly enriched in the blood, but was also increased in the tissue compartments. All compartments were characterized by strong enrichment of pro-inflammatory Mo (PBMC Myeloid Cluster 6, Lung Myeloid Clusters 3 and 6, and BAL Myeloid Cluster 7), although A3 exhibited some differences in these populations compared to A1 and A2. Additionally, this comparison suggested enrichment of AMs in all three compartments; however, upon examination of the specific overlapping gene transcripts, the observed enrichment in blood A1 was primarily related to the presence of non-AM-specific myeloid genes. Finally, numerous common activators and functions of PPI-derived clusters were enriched uniformly across A1, A2, and A3, providing further evidence for pro-inflammatory activity of myeloid cell populations in COVID-19 blood and tissue compartments.

We used trajectory analysis to understand potential transitions of Mo/MΦ in various tissue compartments. We based this on the normalized 196 myeloid-cell specific genes, as well as 425 additional normalized genes that could be important in Mo/myeloid/MΦ cell differentiation, reflective of chemotaxis, IFN, and metabolism genes (**Table S6**). This analysis suggested a branch point of differentiation of Mo/MΦ between blood and lung, with some blood Mo/MΦ differentiating directly to airway cells and others to lung cells in a more protracted manner as indicated by pseudotime (**Figure 5G**).

### Analysis of the biologic activities of myeloid subpopulations

To focus on functional distinctions among the co-expressed myeloid populations in the blood, lung, and airway compartments (A1-A3), we utilized linear regression analyses between GSVA scores for A1-A3 and scores for metabolic, functional, and signaling pathways (**Figure 6, Figure S4**). Blood A1 was significantly correlated with glycolysis, the NLRP3 inflammasome, and the classical and lectin-induced complement pathways. In lung A2, there were no significant correlations detected with metabolism, but this population was significantly correlated with the NLRP3 inflammasome and the alternative complement pathway. Finally, airway A3 was positively correlated with OXPHOS, the classical complement pathway, and TNF signaling and negatively correlated with apoptosis. Overall, these results delineated the heterogeneity in metabolic and inflammatory pathways among myeloid cells enriched in the blood, lung, and airway of COVID-19 patients.

**Figure 6.**
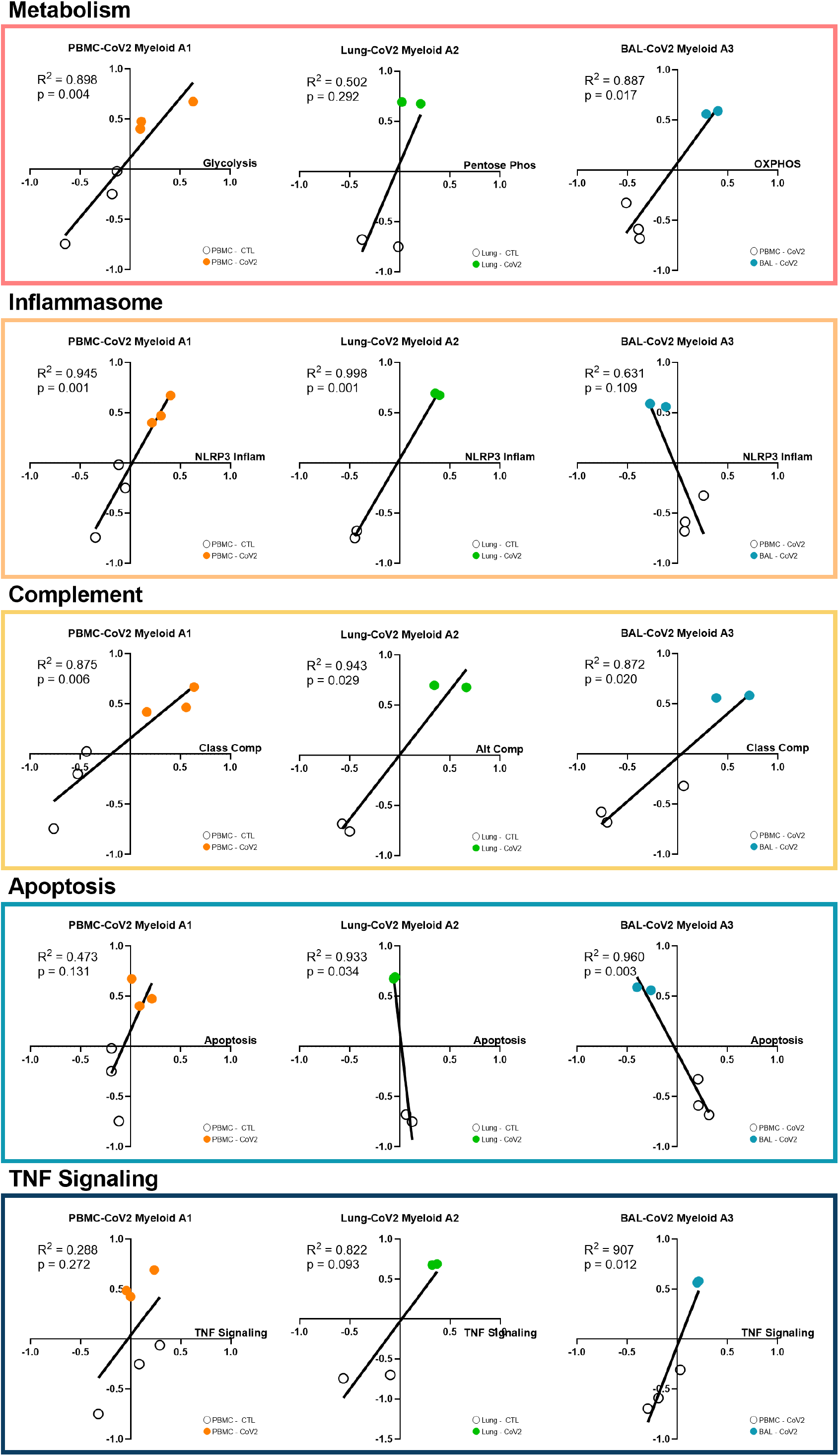
Analysis of biological activities of myeloid subpopulations. Linear regression between GSVA scores for each of the tissue-specific myeloid populations (A1-A3) and metabolism, NLRP3 Inflammasome, complement, apoptosis, and TNF signaling.

### Pathway and upstream regulator analysis inform tissue-specific drug discovery for treatment of COVID-19

To understand the biology of SARS-CoV2-infected patients in greater detail, we conducted pathway analysis on DEGs from the 3 compartments using IPA canonical signaling pathway and upstream regulator (UPR) analysis functions **(Figure 7)**. In general, IFN signaling, the inflammasome, and other components of anti-viral, innate immunity were reflected by disease state gene expression profiles compared to healthy controls **(Figure 7A)**. In addition, metabolic pathways including OXPHOS and glycolysis were significantly increased in the blood of COVID-19 patients compared to controls.

**Figure 7.**
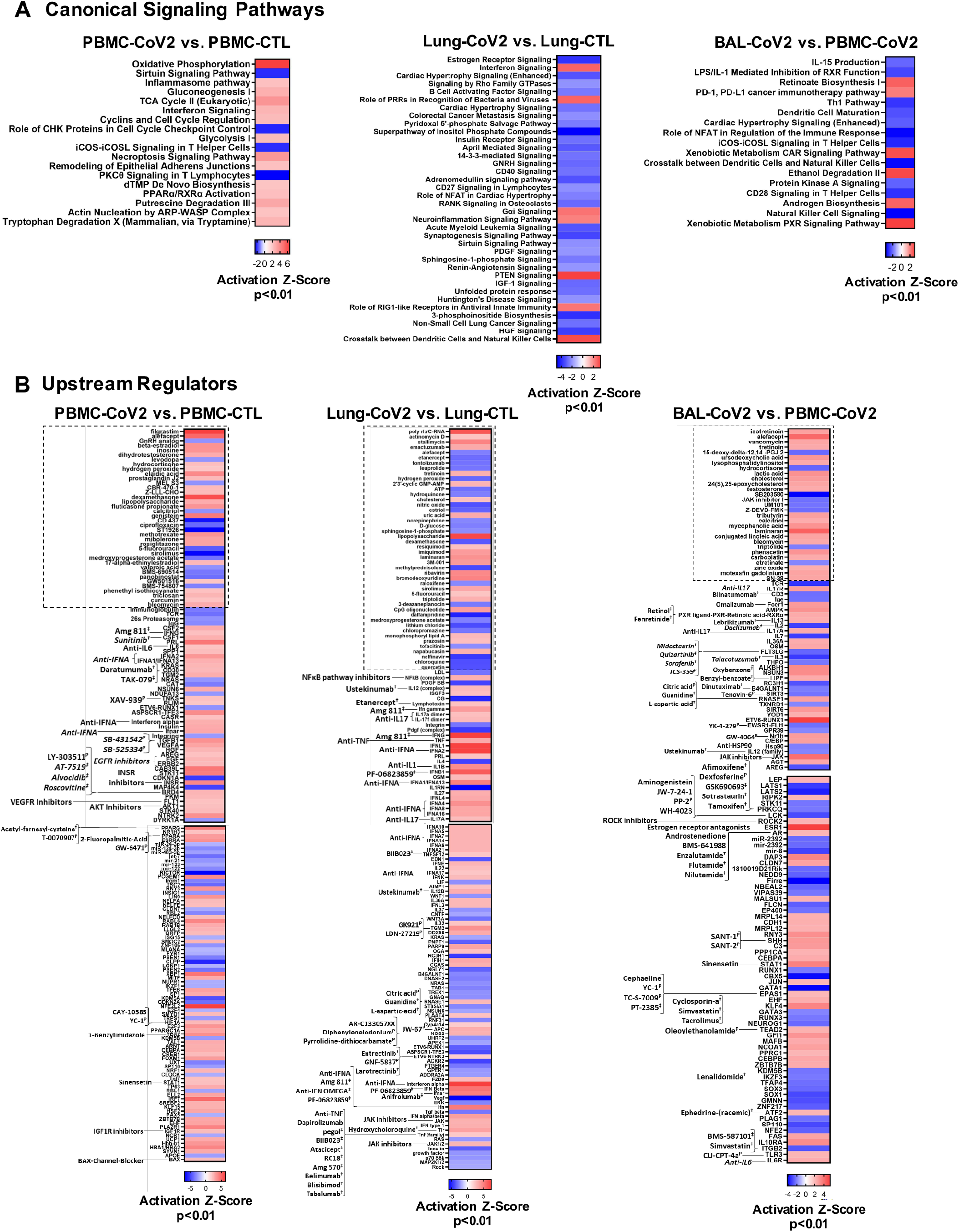
Pathway Analysis of SARS-CoV-2 blood, lung, and airway. DEGs from each SARS-CoV-2 blood or tissue pairwise comparison were uploaded into IPA and canonical signaling pathway (**A**) and upstream regulator (**B**) analyses were performed. Heatmaps represent significant results by Activation Z-Score ≥2 and overlap p-value<0.01. The boxes with the dotted outline separate drugs that were predicted as upstream regulators from pathway molecules and complexes. The remaining, significant upstream regulators were matched with drugs with known antagonistic targeting mechanisms. Drugs in italicized text target the matched upstream regulator by an indirect targeting mechanism. The top 150 UPRs in the lung are shown in (**B**) and the remaining are in Figure S5. ^†^: FDA-approved ^‡^: Drug in development/clinical trials ^P^: Preclinical

UPRs predicted to drive the responses in each compartment indicated uniform involvement of inflammatory cytokines, with Type I IFN regulation dominant in the SARS-CoV2-infected lung **(Figure 7B)**. Notable UPRs of COVID-19 blood included IFNA, IFNG, multiple growth factors and ligands, HIF1A, CSF1 and CSF2. Evidence of inflammatory cytokine signaling by IL17 and IL36A was predicted in COVID-19 lung and airway compartments. Whereas the airway DEG profile indicated regulation by both inflammatory and inhibitory cytokines, the COVID-19 lung UPRs were markedly inflammatory, including, NFκB, IL12, TNF, IL1B, and multiple Type I IFNs. These proinflammatory drivers were consistent in each individual lung which we analyzed separately because of the apparent heterogeneity between the lung samples **(Figure S5).**

IPA analysis was also employed to predict drugs that might interfere with COVID-19 inflammation (**Figure 7B, Table S7**). Of note, neutralizers of IL17, IL6, IL1, IFNA, IFNG, and TNF were predicted as antagonists of COVID-19 biology. Corticosteroids were predicted to revert the gene expression profile in the SARS-CoV-2-infected lung, but were predicted as UPRs of COVID-19 blood, which may indicate that the patients from whom blood was collected had been treated with corticosteroids rather than indicating that these agents were driving disease pathology. Chloroquine (CQ) and hydroxychloroquine (HCQ) were additionally predicted to revert the COVID-19 transcription profile in the lung, which may point to their potential utility as treatment options. A number of drugs matched to unique targetable pathways in the lung, including NFκB pathway inhibitors, antagonists of NOS2, and neutralizers of the TNF family; however, some drugs also targeted pathways shared by both the lung and airway, including anti-RNASE1 compounds and JAK inhibitors. In the BAL-CoV2 vs. PBMC-CoV2 IPA comparison, several drugs were matched to UPRs with a negative Z-score, which provided additional therapeutic options directed towards the blood of patients with COVID-19, given that downregulated, or inhibited UPRs are molecules that could revert the BAL-CoV2 gene signature to that of the PBMC-CoV2 gene signature. As such, several possible therapeutics arose from this analysis to target COVID-19 blood, including ustekinumab, targeting the IL12/23 signaling pathway, and cyclosporine, tacrolimus, and lenalidomide, all of which have immunosuppressive effects. In addition, IGF1R inhibitors, EGFR inhibitors, VEGFR inhibitors, and AKT inhibitors were among the compounds predicted to target COVID-19 PBMCs. No specific drugs were predicted to target all three tissue compartments, but each compartment was driven by inflammatory cytokines.

Another means to predict possible drug targets is by employing connectivity scoring with drug-related gene expression profiles using the perturbagen CMAP database within CLUE (**Table S7, see Methods**). Although CLUE-predicted drugs tended to differ from those predicted by IPA or those matched to IPA-predicted UPRs, there were some overlapping mechanisms, including inhibition of AKT, angiogenesis, CDK, EGFR, FLT3, HSP, JAK, and mTOR. IPA-predicted drugs that were unique from connectivity-predicted drugs tended to capture more cytokine and lymphocyte biology, including inhibitors of IL1, IL6, IL17, TNF, type I and II interferon, CD40LG, CD38, and CD19, among other cytokines and immune cell-specific markers. Overall, our gene expression-based analysis of SARS-CoV2-infected blood and tissue compartments indicated several existing treatment options that could be considered as candidates to treat COVID-19.

## Discussion

Multiple orthogonal bioinformatics approaches were employed to analyze DEG profiles from the blood, lung, and airway of COVID-19 patients and revealed the dynamic nature of the inflammatory response to SARS-CoV-2 and possible points of therapeutic intervention. In the blood, we saw evidence of myeloid cell activation, lymphopenia, and elevation of plasma cells, as has been shown by both standard cell counts, flow cytometry and gene expression analysis (Tay et al., 2020). In the lungs, we found increased gene signatures of additional myeloid cell types including granulocytes, infiltrating inflammatory Mo, and AMs as well as the presence of non-activated T, B, and NK cells. Furthermore, inflammation in the lung tissue was enhanced by the greater presence of IFNs and more pro-inflammatory cytokines than observed in the blood. Finally, in the airway we found evidence of blood and AM-derived inflammatory and regulatory MΦs, and non-hematopoietic lung tissue cells accompanied by expression of SARS-CoV-2 receptors, and alarmins, indicative of viral infection and damage to the lung and consistent with previous reports of detection of SARS-CoV2 in BAL fluid (Corley et al., 2020; Xiong et al., 2020). Together these findings suggest a systemic, but compartmentalized immune/inflammatory response with specific signs of cellular activation in blood, lung and airway. This has informed a more comprehensive and integrated model of the nature of the local and systemic host response to SARS-CoV2.

The predominant populations of immune cells we found to be enriched and activated in COVID-19 patients were myeloid cells and, in particular, subsets of inflammatory Mo and MΦs, which differed between the blood, lung, and airway compartments. In the peripheral blood, we found significant enrichment of Mo, including classically activated inflammatory M1 MΦs as well as a CD33^+^ myeloid subset, which appeared to be an M2 population reminiscent of previously characterized IFN-activated MΦs, AMs, and MDSCs, indicative of a potential regulatory population induced by stimuli arising from the SARS-CoV2-infected lung. Myeloid cells enriched in the blood of COVID-19 patients were also highly correlated with gene signatures of metabolic pathways (Glycolysis, Pentose Phosphate Pathway, and TCA cycle) indicative of pro-inflammatory M1 MΦs (Viola et al., 2019).

The lung tissue was enriched in gene signatures of Mo/MΦs as well as other myeloid cells including neutrophils and LDGs. The role of neutrophils and LDGs in COVID-19 pathogenesis has been poorly characterized, although increases in blood neutrophils have been found to be associated with poor disease outcome in COVID-19 patients and some have proposed that the formation of neutrophil extracellular traps (NETs) contributes to increased risk of death from SARS-CoV2 infection (Barnes et al., 2020; Huang et al., 2020; Thierry and Roch, 2020). Although LDGs have not been previously reported in the COVID-19 lung, in comparison to neutrophils, they exhibit an enhanced capacity to produce Type I IFNs and form NETs and therefore, may have an even greater impact on disease progression (Carmona-Rivera and Kaplan, 2013).

Mo/MΦ subsets in the lung of COVID-19 patients were characterized as infiltrating inflammatory Mo and activated AMs, which exhibited a mixed metabolic status suggestive of different states of activation. Infiltrating Mo from the peripheral blood appeared to be further activated in the lung tissue as evidenced by enhanced expression of markers of highly inflammatory Mo previously characterized in severe COVID-19 cases (Liao et al., 2020). The AM population enriched in COVID lung tissue clustered with genes involved in the coagulation system, which is consistent with observations of procoagulant AM activity in COVID-19 and in ARDS (Tipping et al., 1988). As pulmonary thrombosis has been associated with poor clinical outcomes in COVID-19 patients, this result suggests that activated AMs in SARS-CoV2-infected lung tissue may be involved in facilitating a pro-thrombotic status, and thereby, contribute to poor disease outcome (Wichmann et al., 2020).

In the airway, we detected gene signatures of various post-activated MΦ subsets including inflammatory M1 MΦs, alternatively activated M2 MΦs, and activated AMs. Expression of myeloid cell genes in the airway also correlated with a signature of oxidative metabolism, which is characteristic of M2 macrophages and typically associated with control of tissue damage (Viola et al., 2019). However, in the context of pulmonary infection, polarization of AMs toward an anti-inflammatory M2 phenotype was found to promote continued inflammation, suggesting that these MΦs may not be effective at resolving anti-viral immunity (Allard et al., 2018).

In addition to myeloid cells, inflammatory mediators from the virally infected lung typically promote migration and activation of NK cells and adaptive immune cells including T and B cells (Newton et al., 2016). We found significant deficiencies in gene signatures of T cells and cytotoxic CD8 and NK cells, consistent with clinical evidence of lymphopenia in the peripheral blood and airway of COVID-19 patients (Chen et al., 2020b; He et al., 2005; Qin et al., 2020; Xu et al., 2020). In contrast to T and NK cells, we observed increased evidence of B cell activation through CD40/CD40L and an increased plasma cell signature in the blood of COVID-19 patients. This result suggests that COVID-19 patients are able to mount an antibody-mediated immune response. However, whether a virus-specific antibody response is beneficial to recovery from SARS-CoV2 infection is unclear (Wu et al., 2020). Low quality, low affinity antibody responses to SARS-CoV have been found to promote lung injury in some patients, although it is unknown if this occurs in SARS-CoV2 infected individuals (Iwasaki and Yang, 2020; Liu et al., 2019).

The contents of the airway as assessed through the BAL fluid, act as a window into events in the alveoli and airways and can be used to understand what is happening in the infected tissue that is separate from the interstitium of the lung (Hamacher et al., 2002; Heron et al., 2012). We found increased enrichment of lung epithelial cells in the airway of COVID-19 patients, suggesting that SARS-CoV2 infection of alveolar cells together with localized inflammation as a result of enhanced myeloid cell infiltration promote significant damage to the alveoli and result in affected cells being sloughed into the airway. Furthermore, the lack of cytotoxic cells and thus, the inability to clear these virus-infected lung epithelial cells in the airway likely accounts for the increased presence of post-activated MΦs and high expression of pro-inflammatory IL-1 family members we observed in the BAL of COVID-19 patients. Importantly, these results suggest that sampling of BAL may provide an important mechanism to evaluate the impact of SARS-CoV2 infection.

DEGs from COVID-19 patients were enriched in IGS, complement pathways, inflammatory cytokines and the inflammasome, which would be expected to activate Mo/MΦ populations in the blood, lung, and airway of COVID-19 patients and initiate a robust and systemic response to infection. In particular, our results support the conclusion that IL-1 family-mediated inflammation plays a critical role in COVID-19 pathogenesis. However, in contrast to previous characterizations of both SARS-CoV and SARS-CoV2 infection, our analysis did not indicate the presence of overwhelming pro-inflammatory cytokine production or “cytokine storm” in COVID-19 patients (Sun et al., 2020; Tay et al., 2020). Rather, our data suggests that the increased numbers, overactivation, and potentially heightened pathogenicity of monocyte/MΦ populations are the main drivers of the dysregulated inflammatory response and resulting tissue damage in COVID-19 patients.

In the absence of proven antiviral treatment and/or a SARS-CoV2-specific vaccine, disease management is reliant upon supportive care and therapeutics capable of limiting the severity of clinical manifestations. Using empiric evidence as a guide, the current approach has been successful in identifying “actionable” points of intervention in an unbiased manner and in spite of formidable patient heterogeneity. Analyses presented here support several recent reports highlighting COVID-19 infection-related increases in inflammatory cytokines, particularly IL6 and TNF, both of which function as predictors of poor prognosis (Pedersen and Ho, 2020; Voiriot et al., 2017), as well as complement activation (Gao et al., 2020; Huang et al., 2020; Mehta et al., 2020). Accordingly, anti-IL6 therapies including sarilumab, tocilizumab and clazakizumab, as well as biologics targeting terminal components of the complement cascade, such as eculizumab and ravulizumab, are in various clinical trial phases for treating COVID-19-associated pneumonia. Candidate TNF blockers such as adalimumab, etanercept and many others, represent additional options for inhibiting deleterious pro-inflammatory signaling. However, most showed patient heterogeneity, suggesting a requirement to identify the specific cytokine profile in each patient in order to offer personalized treatment. Our analyses also point to the likely involvement of pro-inflammatory IL1 family members especially in the lung, suggesting anti-IL1 family interventions, including canakinumab and anakinra, may be effective in preventing acute lung injury.

This analysis also establishes the predominance of inflammatory Mo/myeloid lineage cells in driving disease pathology and suggests therapies effective at blocking myeloid cell recruitment or forcing repolarization may prevent disease progression. CCL5 (RANTES) is a potent leukocyte chemoattractant that interacts with multiple receptors, including CCR1 (upregulated in the blood, lung and airway), and CCR5 (upregulated in the airway). Disruption of the CCR5-CCL5 axis was recently tested using the CCR5 neutralizing monoclonal antibody leronlimab in a small compassionate use trial with promising preliminary results (Patterson et al., 2020).

It has also been observed that COVID-19 may predispose patients to thromboembolic disease (Klok et al., 2020; Middeldorp et al., 2020). Indeed, the gene expression analyses presented here showing altered expression of coagulation factors and fibrinogen genes suggests dysfunction within the intrinsic clotting pathway. These findings, together with evidence of excessive inflammation, complement activation and the involvement of LDGs in lung inflammation, may contribute to the systemic coagulation underlying the remarkably high incidence of thrombotic complications observed in severely ill patients, thereby reinforcing recommendations to apply pharmacological anti-thrombotic medications.

Finally, there has been much recent discussion concerning the use of anti-rheumatic drugs for managing COVID-19. In fact, CQ was one compound predicted as a UPR with potential phenotype-reversing properties. In vitro experiments examining the anti-viral properties of CQ and its derivative HCQ were effective in limiting viral load; however, the efficacy of these drugs in clinical trials has been less clear (Geleris et al., 2020). Questions about drug timing, dosage and adverse events have all called into question the use of these drugs for COVID-19 patients. Despite a recent report showing no negative connectivity between the gene signatures of SARS-CoV2 infection and HCQ treatment (Corley et al., 2020), IPA predicted a role of anti-malarials as limiting the function of intracellular TLRs in the lung and also as a direct negative UPR of gene expression abnormalities in the lung, suggesting a role in controlling COVID-19 inflammation and not viral replication. Further clinical testing may be necessary to test this possible utility.

By comparing the transcriptomic profile of the blood, lung, and airway in COVID-19 patients, a model of the systemic pathogenic response to SARS-CoV2 infection has emerged (Graphical abstract). SARS-CoV2 infection leads to systemic Mo/MΦ activation, likely as a result of the release of pro-inflammatory mediators from infected cells. Infiltration of immune cells into the lung tissue and alveolus, in particular, neutrophils, LDGs, and pathogenic Mo/MΦ populations promotes a cycle of inflammatory mediator release and further myeloid cell activation, which exacerbates inflammation in the lung and leads to tissue damage. The local release of complement components and clotting factors by infiltrating Mo/MΦ may contribute to both inflammation and thrombotic events. As disease progresses, increased infiltration of pro-inflammatory immune cells, release of inflammatory mediators, and damage to the infected alveolus is reflected by the presence of Mo/MΦ cells and lung epithelial cells in the airway (Heron et al., 2012). Furthermore, evidence of both Mo-derived inflammatory MΦs and AMs in the airway compartment suggests that myeloid cell populations from both the blood and the lung tissue are present in the BAL. The accumulation of virus-infected cells and release of alarmins in the airway may not only reflect ongoing infection, but also promote inflammation and prevent resolution of the infection and foster the continuation of the innate immune response. Therefore, sampling the BAL fluid seems to be an effective strategy to monitor tissue inflammation and damage in COVID-19 patients.

As SARS-CoV2 continues to propagate, viral clearance is impaired by a lack of cytotoxic CD8 T cells and NK cells. This is consistent with MAS occurring in other settings, in which defects in cytotoxic activity of CD8 T cells and NK cells result in enhanced innate immune cell activation and intensified production of pro-inflammatory cytokines, many of which were also expressed in COVID-19 patients (Crayne et al., 2019; McGonagle et al., 2020). Thus, we propose that the lack of activated CD8 T cells and NK cells and subsequent failure to clear virus-infected cells, is a major contributor to the MΦ-driven pathologic response to SARS-CoV2 observed in COVID-19 patients.

In order to develop a model of SARS-CoV2 infection, we have utilized multiple orthogonal approaches to analyze gene expression from COVID-19 patients, but also acknowledge the limitations of this data. For example, we were only able to analyze 2-3 samples per experimental condition and this limited the statistical power of each of our bioinformatics techniques. In addition, the low number of samples meant that heterogeneity among patients in any given cohort had an increased impact on the overall outcome. One possible reason for this intra-cohort heterogeneity is that the patients may have exhibited varying levels of disease severity and, unfortunately, we did not have access to clinical information. These points highlight the need for additional studies on more patients, preferably accounting for differences in demographic information and disease status, in order to increase the power of downstream analyses.

In conclusion, transcriptomic analysis has contributed critical insights into the pathogenesis of COVID-19. Diffuse Mo/MΦ activation is the likely primary driver of clinical pathology. Therefore, this work provides a rationale for placing greater focus on the detrimental effects of exaggerated activation of pathogenic Mo/MΦs and for targeting these populations as an effective treatment strategy for COVID-19 patients.

## Methods

### Read quality, trimming, mapping and summarization

Publicly available data sets used in this study are listed in **Table S1**. RNA-seq data were processed using a consistent workflow using FASTQC, Trimmomatic, STAR, Sambamba, and featureCounts. As described below SRA files were downloaded and converted into FASTQ format using SRA toolkit. Read ends and adapters were trimmed with Trimmomatic (v0.38) using a sliding window, ilmnclip, and headcrop filters. Both datasets were head cropped at 6bp and adapters were removed before read alignment. Reads were mapped to the human reference genome hg38 using STAR, and the .sam files were converted to sorted .bam files using Sambamba. Read counts were summarized using the featureCounts function of the Subread package (v1.61.)

The RNA-seq tools are all free, open source programs available at the following web addresses

SRA toolkit - https://github.com/ncbi/sra-tools

FastQC - https://www.bioinformatics.babraham.ac.uk/projects/fastqc/

Trimmomatic - http://www.usadellab.org/cms/?page=trimmomatic

STAR - https://github.com/alexdobin/STAR

http://labshare.cshl.edu/shares/gingeraslab/www-data/dobin/STAR/STAR.posix/doc/STARmanual.pdf

Sambamba - http://lomereiter.github.io/sambamba/

FeatureCounts - http://subread.sourceforge.net/

### Differential gene expression and gene set enrichment analysis

The DESeq2 workflow was used for differential expression analysis. Comparisons were made between control PBMCs and PBMCs from COVID-19 patients (PBMC-CTL vs PBMC-CoV2) and control lung tissue and lung tissue from COVID-19 patients (Lung-CTL vs Lung-CoV2). Since no corresponding control BAL samples were available for the COVID-19 BAL samples, we compared BAL samples from COVID-19 patients to COVID-19 PBMC (PBMC-CoV2 vs BAL-CoV2). This was possible because these samples were analyzed on the same platform, run at the same time, and it was done understanding the limitations of this analysis. We also compared normal BAL to BAL of asthmatic individuals to identify genes unrelated to COVID-19 (PRJNA434133)

Two technical replicates were included for BAL cohort, and 4 technical replicates were included for postmortem lung samples. The replicates were collapsed and averaged into one using collapsereplicates function from DESeq2 package. The genes with low expression (i.e genes with very few reads) were removed by filtration. The filtered raw counts were normalized using the DESeq method and differentially expressed genes were determined by FDR < 0.2 (Anders and Huber, 2010). Counts were then log2 transformed and used for downstream analyses (**Table S2**).

### Gene Set Variation Analysis (GSVA)

The GSVA (Hänzelmann et al., 2013) (V1.25.0) software package is an open source package available from R/Bioconductor and was used as a non-parametric, unsupervised method for estimating the variation of pre-defined gene sets in patient and control samples of microarray and RNA-seq expression data sets (www.bioconductor.org/packages/release/bioc/html/GSVA.html). The inputs for the GSVA algorithm were a gene expression matrix of log_2_ expression values for pre-defined gene sets (**Table S3**). All genes within a gene set were evaluated if the interquartile range (IQR) of their expression across the samples was greater than 0. Enrichment scores (GSVA scores) were calculated non-parametrically using a Kolmogorov Smirnoff (KS)-like random walk statistic and a negative value for a particular sample and gene set, indicating that the gene set has a lower expression than the same gene set with a positive value. The enrichment scores (ES) were the largest positive and negative random walk deviations from zero, respectively, for a particular sample and gene set. The positive and negative ES for a particular gene set depend on the expression levels of the genes that form the pre-defined gene set. GSVA calculates enrichment scores using the log_2_ expression values for a group of genes in each SARS-CoV2 patient and healthy control and normalizes these scores between −1 (no enrichment) and +1 (enriched). Welch’s t-test was used to calculate the significance of the gene sets between the cohorts. Significant enrichment of gene sets was determined by p-value < 0.05.

### Derivation of GSVA Gene Sets

All input gene sets used for GSVA analysis can be found in **Table S3**. Cellular and inflammatory modules (Catalina et al. submitted manuscript) and IFN-induced gene sets were previously derived (Catalina et al., 2019). Additional inflammatory pathways (NLRP3 Inflammasome, Classical Complement, Alternative Complement, and Lectin-induced Complement were curated from the Molecular Signatures Database. Other signatures were derived from various publications (Culemann et al., 2019; Kotliarov et al., 2020; Kuo et al., 2019; Liao et al., 2020; Martinez et al., 2006; Reyfman et al., 2019).

Additional hematopoietic cellular gene signatures (monocyte, myeloid, and neutrophil) were derived from I-Scope™, a tool developed to identify immune cell specific genes in big data gene expression analyses. Non-hematopoietic fibroblast and lung cell gene sets were derived from T-Scope™, a tool developed to identify genes specific for 45 non-hematopoietic cell types or tissues in big gene expression datasets. The T-Scope™ database contains 1,234 transcripts derived initially from 10,000 tissue enriched and 8,000 cell line enriched genes listed in the Human Protein Atlas. From the list of 18,000 potential tissue or cell specific genes, housekeeping genes and genes differentially expressed in 40 hematopoietic cell datasets were removed. The final gene lists were checked against available single cell analyses to confirm cellular specificity.

Nine single-cell RNA-seq lung cell populations (AT1, AT2, Ciliated, Club, Endothelial, Fibroblasts, Immuno Monocytes, Immuno T Cells, and Lymphatic Endothelium) were downloaded from the Eils Lung Tissues set (Lukassen et al., 2020) accessed by the UC Santa Cruz Genome Browser (https://eils-lung.cells.ucsc.edu). Genes occurring in more than one cell type were removed. Additionally, genes known to be expressed by immune cells were removed. The Eils Lung Tissues set Immuno Monocyte, Immuno T Cell, Fibroblast, and Lymphatic Endothelium categories were not employed in further analyses.

Apoptosis and NFkB gene signatures were derived and modified from Ingenuity® Pathway Analysis pathways Apoptosis Signaling and NFkB Signaling. ROS-protection was derived from Biologically Informed Gene-Clustering (BIG-C®).

### Network analysis and visualization

Visualization of protein-protein interaction and relationships between genes within datasets was done using Cytoscape (version 3.6.1) software (**Table S4**). Briefly, STRING (version 1.3.2) generated networks were imported into Cytoscape (version 3.6.1) and partitioned with MCODE via the clusterMaker2 (version 1.2.1) plugin. For PPIs in **Figure 4A-C**, STRING settings were adjusted to high confidence (0.7), for PPIs in **Figure 4D-F**, settings were relaxed to medium confidence (0.4). All PPIs were generated without the neighborhood or textmining features. For some PPIs, the average interaction strength using STRING-based cumulative interaction scores was used to determine the strength of interaction between clusters.

### Functional and cellular enrichment analysis

Functional enrichment of clusters was performed using Biologically Informed Gene-Clustering (BIG-C®), which was developed to understand the potential biological meaning of large lists of genes (Labonte et al., 2018). Genes are clustered into 53 categories based on their most likely biological function and/or cellular localization based on information from multiple on-line tools and databases including UniProtKB/Swiss-Prot, GO terms, KEGG Pathways, MGI database, KEGG pathways, NCBI PubMed, and the Interactome. Hematopoietic cellular enrichment was performed using I-Scope™, a tool developed to identify immune cell specific genes in big data gene expression analyses.

Statistically significant enriched types of cell types in DEGs are determined by Fisher’s Exact test overlap p-value and then determining an Odds Ratio of enrichment.

### Derivation of co-expressed myeloid subpopulations in each compartment

Co-expression analyses were conducted in R. Sample (control and patient) log2 expression values for each gene of the 221 identified monocyte/myeloid cell genes in were analyzed for their Pearson correlation coefficient in each tissue compartment (blood, lung, and airway) using the Cor function. Of note, only 196 of 221 genes had changes in gene expression in at least one tissue by RNA-seq. Pearson correlations for these 196 genes were hierarchically clustered by their Euclidian distance into 2 clusters (k=2) using the heatmap.2 function in R. This resulted in 2 Mo/myeloid co-expressed clusters in each compartment corresponding to increased and decreased gene sets. The upregulated co-expressed genes were used to define the A1, A2, and A3 myeloid subpopulations from the blood, lung, and airway compartments, respectively (**Table S5**). The co-expressed myeloid populations in each compartment (A1-A3) were then evaluated for enrichment by GSVA.

### Inter-compartment myeloid gene comparisons

To compare relative expression of the 196 myeloid-specific genes among compartments, HTS filtered log2 expression values for each gene were normalized to the average expression of *FCGR1A*, *FCGR2A*, and *FCGR2C* in each sample. Welch’s t-test was used to calculate the significant differences in normalized gene expression between cohorts. Effect sizes were computed between cohorts using the cohen.d function with Hedges’ correction in R.

### Monocle

Trajectory analyses were performed with Monocle (Qiu et al., 2017a, 2017b; Trapnell et al., 2014) version 2.14.0 in R. Gene expression values for 621 genes related to myeloid cell differentiation and function including cell surface and secreted markers, M1 and M2 markers, metabolism, and IFN genes were selected as a curated input list (**Table S5**). The HTS filtered log2 expression values for each of these genes in each sample for each tissue type (PBMC-CoV2, Lung-CoV2, and BAL-CoV2) was normalized by the average log2 expression of *FCGR1A*, *FCGR2A*, and *FCGR2C* in that particular sample as described above. Normalized expression of these genes was used as the input expression data for Monocle. The CellDataSet was created with parameters of lowerDetectionLimit = 0.01 and expressionFamily = uninormal(). Dimensions were reduced using the DDRTree method, and the max_components parameter was set to 2. Cell state was ordered based upon the state corresponding to PBMC-CoV2.

### Linear Regression Analysis

Simple linear regression between calculated myeloid subpopulation A1, A2, and A3 GSVA scores and biological functions or signaling pathway GSVA scores was performed in GraphPad Prism Version 8.4.2. In all analyses where pathway genes (e.g. classical complement) were also myeloid cell genes, these genes were removed from the myeloid GSVA score for that comparison and kept in the pathway GSVA score. For each regression analysis, the Goodness of Fit is displayed as the R squared value and the p-value testing the significant of the slope is displayed. All p-values are displayed with 3 digits and rounded-up unless rounding changes significance.

### Ingenuity ® Pathway Analysis

The canonical pathway and upstream regulator functions of IPA core expression analysis tool (Qiagen) were used to interrogate DEG lists. Canonical pathways and upstream regulators were considered significant if |Activation Z-Score|≥2 and overlap p-value<0.01. Chemical reagents, chemical toxicants, and endogenous non-mammalian ligands were culled from all upstream regulator analyses.

### Drug-Target Matching

IPA-predicted upstream regulators were annotated with respective targeting drugs and compounds to elucidate potential useful therapies in SARS-CoV2. Drugs targeting gene products of interest by both direct and indirect targeting mechanisms were sourced by Combined Lupus Treatment Scoring (CoLTS)-scored drugs (Grammer et al., 2016), the Connectivity Map via the drug repurposing tool, DrugBank, and literature mining. Similar methods were employed to determine information about drugs and compounds, including mechanism of action and stage of clinical development. The drug repurposing tool was accessed at clue.io/repurposing-app.

### Analysis of COVID-19 PBMC, Lung, and BAL DEG profiles via CLUE

DEGs from PBMC-CoV2 vs. PBMC-CTL, Lung-CoV2 vs. Lung-CTL, and BAL-CoV2 vs. PBMC-CoV2 were used as input for the CMaP and LINCS Unified Environment (CLUE) cloud-based connectivity map analysis platform (https://clue.io/connectopedia/). Top upregulated and downregulated DEGs from each signature as determined by magnitude of log_2_ fold change were sequentially entered into CLUE until 150 of each were accepted for analysis to determine drugs, compounds, small molecules, and other perturbagens that mimic or oppose the uploaded COVID-19 gene expression signatures. Resultant drugs and compounds with negative connectivity scores in the [−75, −100] range were analyzed to include results with high confidence of antagonizing COVID-19 gene expression profiles.

## Supplemental Figures

**Figure S1.** Metaclusters identify downregulated cell populations and functional gene clusters in SARS-CoV2-infected tissues.

**Figure S2.** Evaluation of published macrophage gene signatures in myeloid-derived clusters from COVID-affected blood, lung and BAL fluid.

**Figure S3.** Heterogeneous expression of monocyte/myeloid cell genes in different tissue compartments.

**Figure S4.** Linear regression analysis reveals significant correlation between myeloid cell populations and additional metabolic and immune signaling pathways.

**Figure S5.** Pathway analysis of SARS-Cov2 lung tissue.

## Supplemental Tables

**Table S1.** Datasets used in study.

**Table S2.** DEGs in COVID-19 blood, lung and airway.

**Table S3.** Gene sets for GSVA analyses.

**Table S4**. MCODE clusters in COVID-19 blood, lung, airway, NHBE epithelial cell line and PPI myeloid sub-clusters.

**Table S5.** Co-expression derived myeloid populations A1, A2, and A3 and their overlaps in the blood, lung, and airway. Input genes for trajectory analysis.

**Table S6.** Enrichment of published macrophage signatures and PPI myeloid sub-clusters in A1, A2, and A3.

**Table S7.** IPA and CLUE-predicted drugs targeting COVID-19 blood, lung, and airway.

