## Supplemental Figures for "Comprehensive Transcriptomic Analysis of COVID-19 Blood, Lung, and Airway"

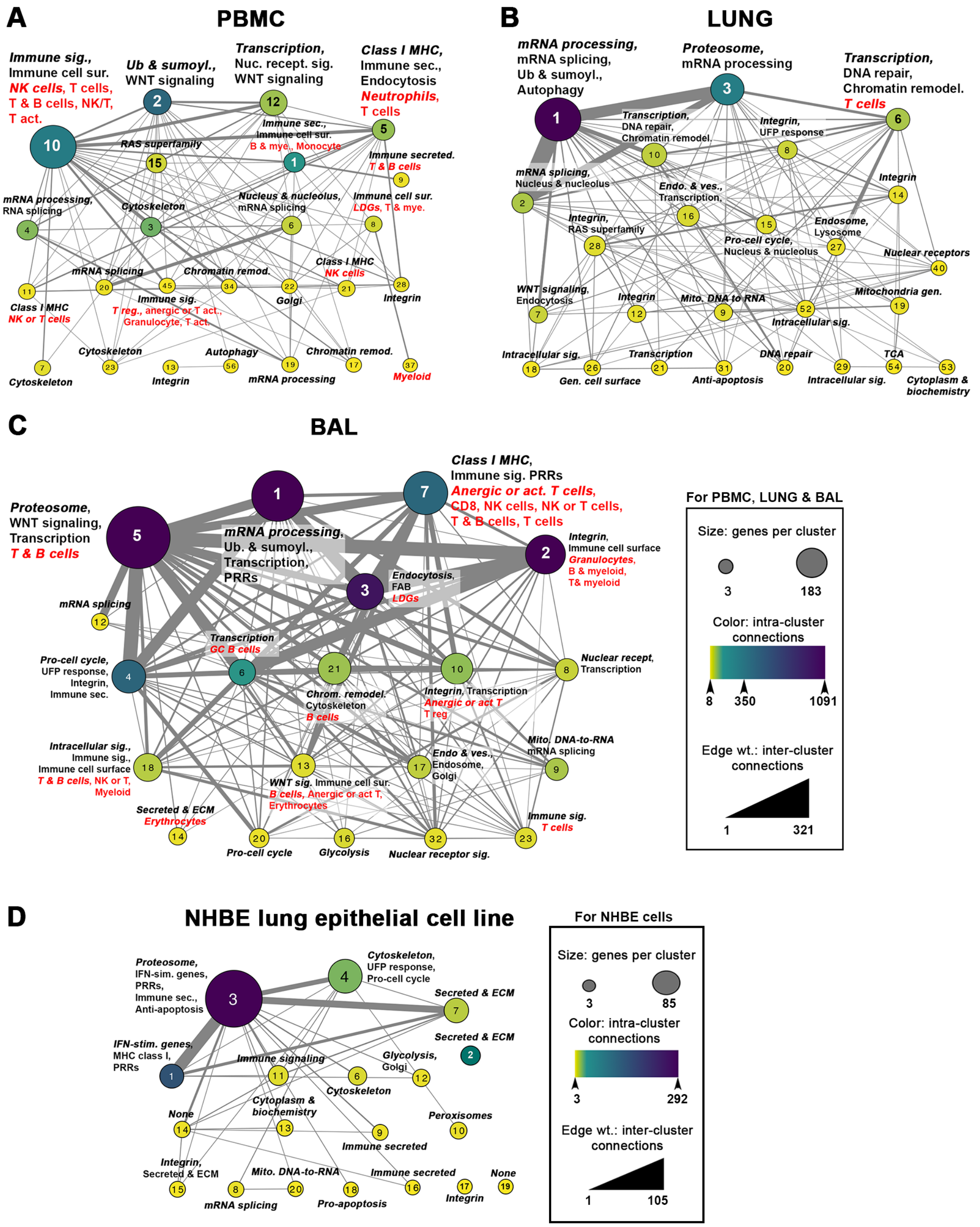


**Figure S1. Metaclusters identify differentially expressed cell populations and functional gene clusters in SARS-CoV2 infected tissues and cell lines.** Down-regulated DE genes from peripheral blood (**A**), lung (**B**) and airway (**C**), and up-regulated DE genes from the NHBE primary lung epithelial cell line (**D**) were used to create metaclusters. Metaclusters were generated based on PPI networks, clustered using MCODE and visualized in Cytoscape. Size indicates the number of genes per cluster, color indicates the number of intra-cluster connections and edge weight indicates the number of inter-cluster connections. Cluster enrichment for biological function and immune cell type was determined by BIG-C and I-Scope, respectively.

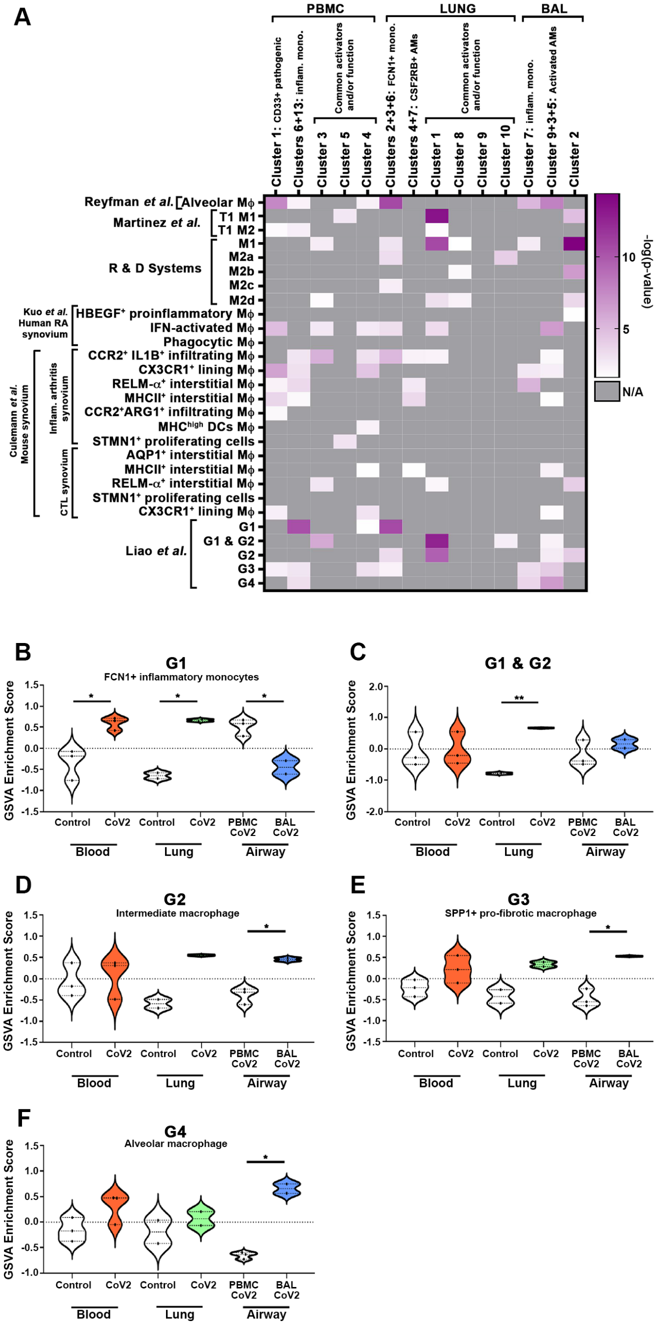


**Figure S2. Evaluation of published macrophage gene signatures in myeloid-derived clusters from COVID-affected blood, lung and BAL fluid.** Previously published macrophage signatures from the indicated sources were compared to myeloid clusters from Figure 3D-F. Heat map depicts signatures with significant overlap (-log(p-value) >1.33) with myeloid clusters from the blood, lung and airway compartments. N/A, non-applicable/non-significant overlap detected. R & D Systems provided signatures for the M1, M2a, M2b, M2c and M2d populations.

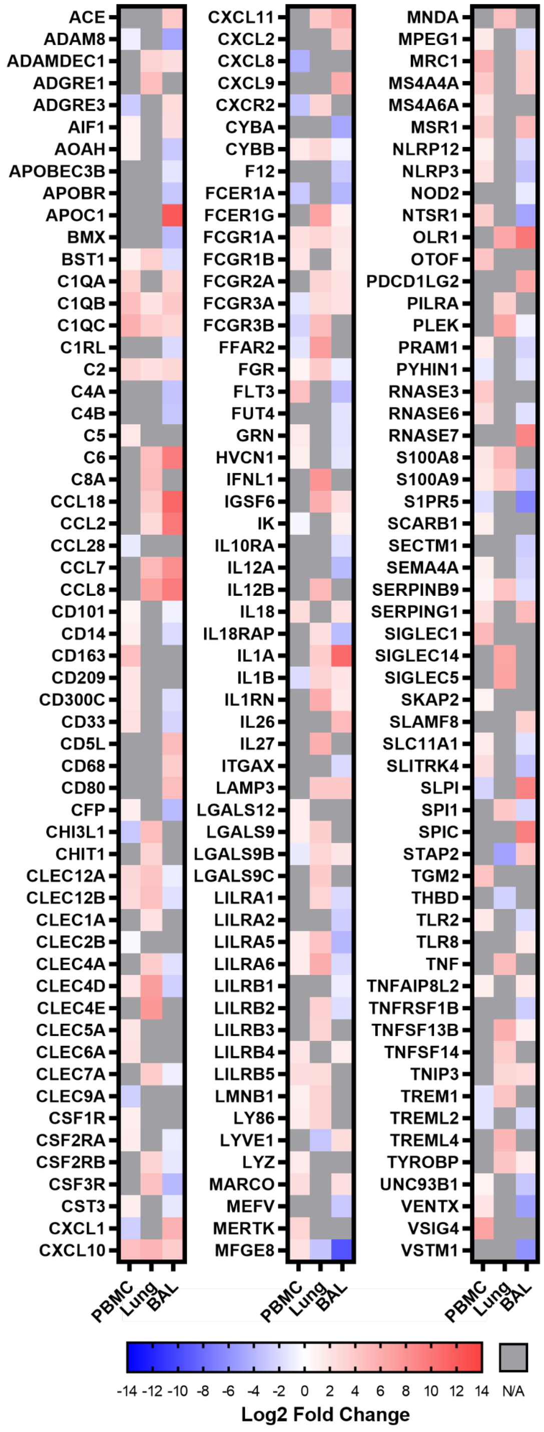


**Figure S3. Heterogeneous expression of monocyte/myeloid cell genes in different CoV2 tissue compartments as compared to control.** Evaluation of differential expression of 171 monocyte/myeloid genes in each compartment reveals shared and disparate expression among the tissues. PBMC represents PBMC-CoV2 to PBMC-CTL. Lung represents Lung-CoV2 to Lung-CTL. BAL represents BAL-CoV2 to PBMC-CoV2. Scale bar presents Log2 Fold Change. N/A represents genes that were not significantly DE at FDR < 0.2.

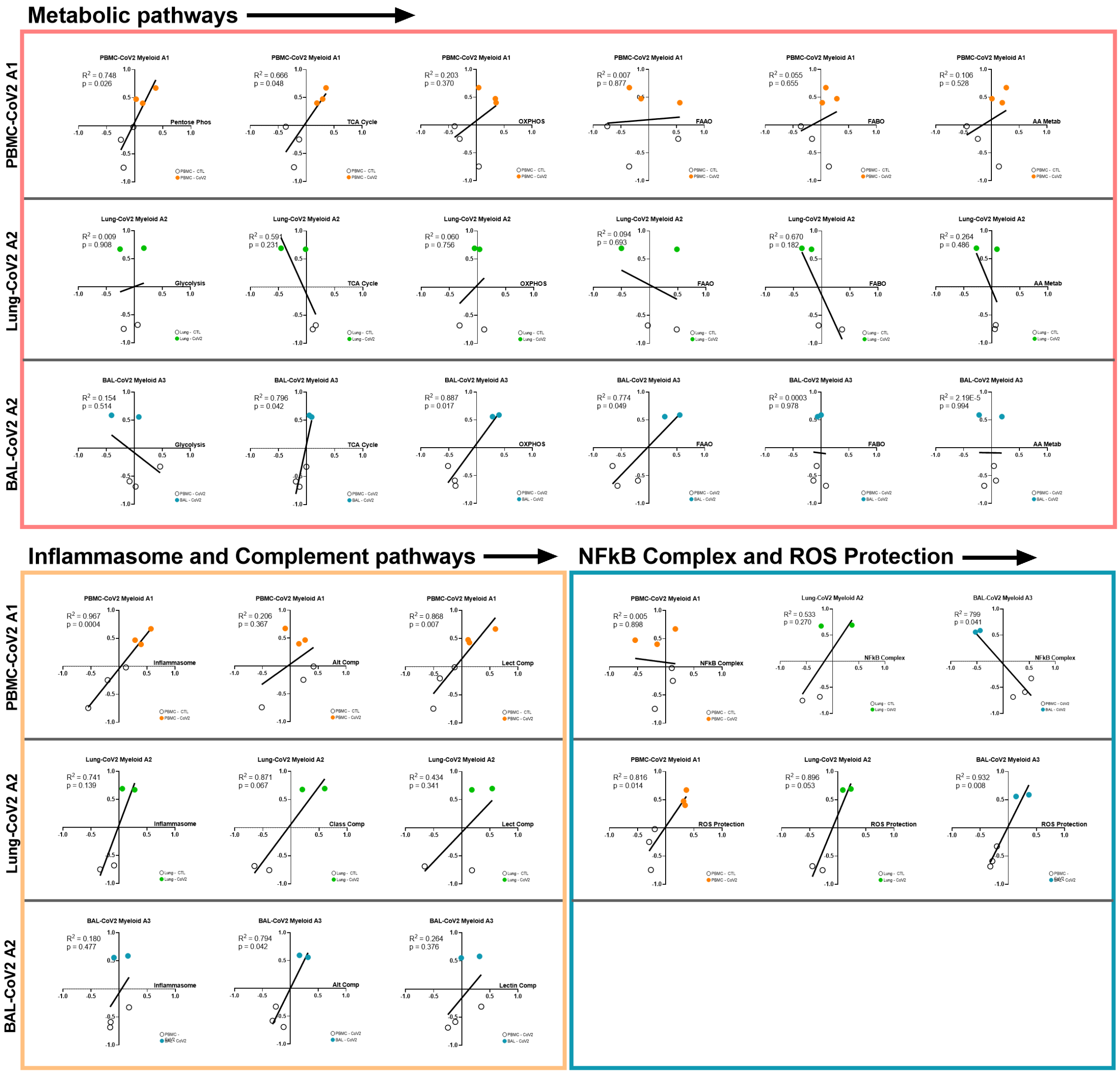


**Figure S4. Analysis of biological activities of myeloid subpopulations.** Linear regression between GSVA scores for each of the tissue-specific myeloid populations and metabolic pathways, inlammasome, complement pathways, NFKB complex signaling and ROS protection.

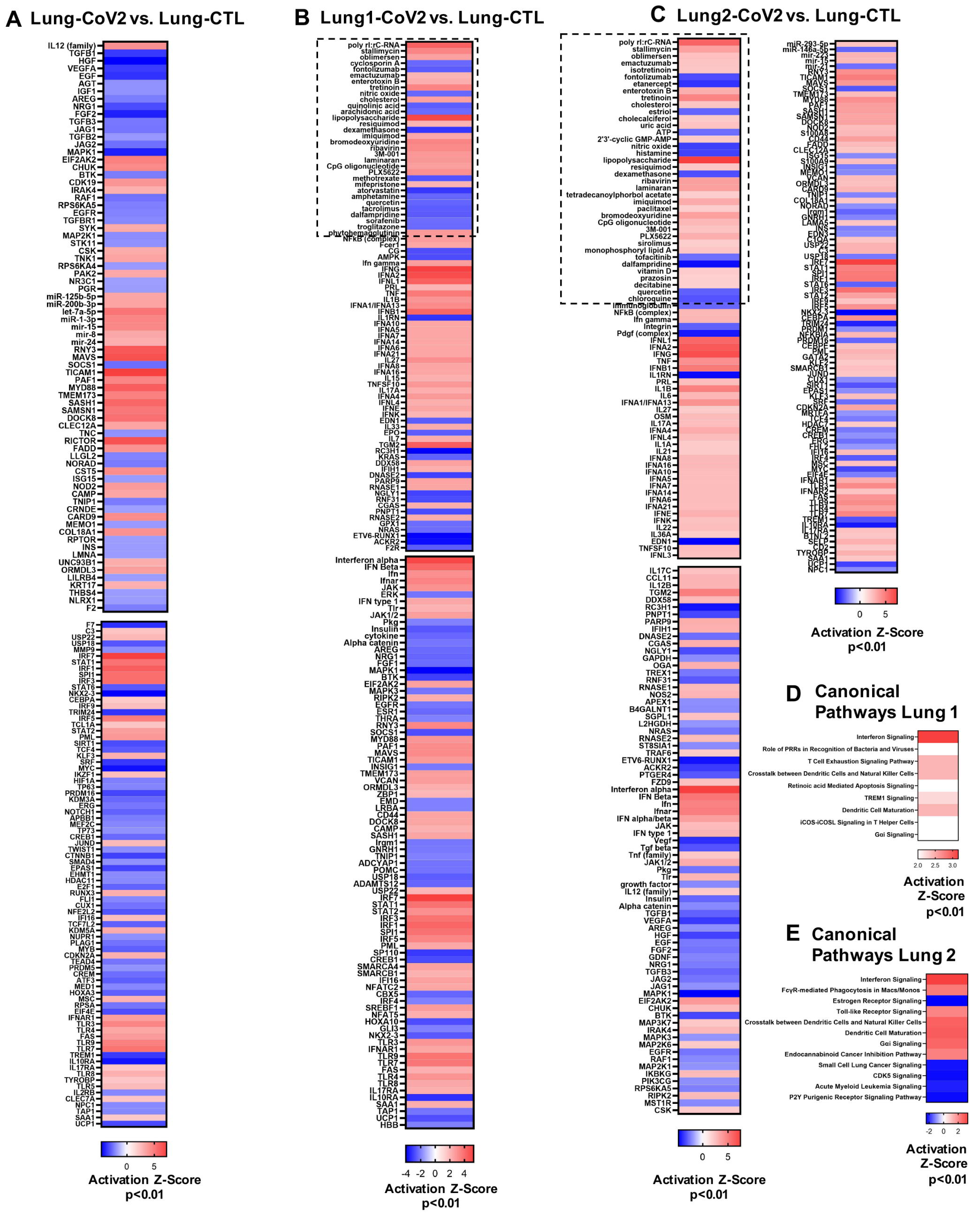


**Figure S5. Pathway analysis of SARS-CoV-2 lung tissue. (A)** Remaining significant upstream regulators operative in SARS-CoV-2 lung tissue predicted by IPA upstream regulator analysis (see Figure 9). Upstream regulator analysis was also conducted on DEGs from each individual COVID-19 lung compared to healthy controls due to observed heterogeneity, with significant results displayed in **(B)** and **(C)**. Chemical reagents, chemical toxicants, and non-mammalian endogenous chemicals were culled from results. The boxes with the dotted outline separate small molecules/drugs/compounds that were predicted as upstream regulators from pathway molecules and complexes. **(D, E)** IPA canonical signaling pathway analysis was conducted on individual COVID-19 lung samples. Pathways and upstream regulators were considered significant by |Activation Z-Score|≥2 and overlap p-value<0.01.
